# Ecophysiology of freshwater Verrucomicrobia inferred from metagenome-assembled genomes

**DOI:** 10.1101/150078

**Authors:** Shaomei He, Sarah LR Stevens, Leong-Keat Chan, Stefan Bertilsson, Tijana Glavina del Rio, Susannah G Tringe, Rex R Malmstrom, Katherine D McMahon

**Author notes:** **Corresponding author** Katherine D McMahon.

## Abstract

Microbes are critical in carbon and nutrient cycling in freshwater ecosystems. Members of the Verrucomicrobia are ubiquitous in such systems, yet their roles and ecophysiology are not well understood. In this study, we recovered 19 Verrucomicrobia draft genomes by sequencing 184 time-series metagenomes from a eutrophic lake and a humic bog that differ in carbon source and nutrient availabilities. These genomes span four of the seven previously defined Verrucomicrobia subdivisions, and greatly expand the known genomic diversity of freshwater Verrucomicrobia. Genome analysis revealed their potential role as (poly)saccharide-degraders in freshwater, uncovered interesting genomic features for this life style, and suggested their adaptation to nutrient availabilities in their environments. Between the two lakes, Verrucomicrobia populations differ significantly in glycoside hydrolase gene abundance and functional profiles, reflecting the autochthonous and terrestrially-derived allochthonous carbon sources of the two ecosystems respectively. Interestingly, a number of genomes recovered from the bog contained gene clusters that potentially encode a novel porin-multiheme cytochrome *c* complex and might be involved in extracellular electron transfer in the anoxic humic-rich environment. Notably, most epilimnion genomes have large numbers of so-called “Planctomycete-specific” cytochrome *c*-containing genes, which exhibited nearly opposite distribution patterns with glycoside hydrolase genes, probably associated with the different environmental oxygen availability and carbohydrate complexity between lakes/layers. Overall, the recovered genomes are a major step towards understanding the role, ecophysiology and distribution of Verrucomicrobia in freshwater.

**IMPORTANCE:** Freshwater Verrucomicrobia are cosmopolitan in lakes and rivers, yet their roles and ecophysiology are not well understood, as cultured freshwater Verrucomicrobia are restricted to one subdivision of this phylum. Here, we greatly expand the known genomic diversity of this freshwater lineage by recovering 19 Verrucomicrobia draft genomes from 184 metagenomes collected from a eutrophic lake and a humic bog across multiple years. Most of these genomes represent first freshwater representatives of several Verrucomicrobia subdivisions. Genomic analysis revealed Verrucomicrobia as potential (poly)saccharide-degraders, and suggested their adaptation to carbon source of different origins in the two contrasting ecosystems. We identified putative extracellular electron transfer genes and so-called “Planctomycete-specific” cytochrome *c*-containing genes, and found their distinct distribution patterns between the lakes/layers. Overall, our analysis greatly advances the understanding of the function, ecophysiology and distribution of freshwater Verrucomicrobia, while highlighting their potential role in freshwater carbon cycling.

## INTRODUCTION

Verrucomicrobia are ubiquitous in freshwater and exhibit a cosmopolitan distribution in lakes and rivers. They are present in up to 90% of lakes (1), with abundances typically between <1% and 6% of total microbial community (2-4), but as high as 19% in a humic lake (5). Yet, in comparison to other freshwater bacterial groups, such as members of the Actinobacteria, Cyanobacteria and Proteobacteria phyla, Verrucomicrobia have received relatively less attention, and their functions and ecophysiology in freshwater are not well understood.

As a phylum, Verrucomicrobia (V) was first proposed relatively recently, in 1997 (6). Together with Planctomycetes (P), Chlamydiae (C), and sister phyla such as Lentisphaerae, they comprise the PVC superphylum. In addition to being cosmopolitan in freshwater, Verrucomicrobia have been found in oceans (7, 8), soil (9, 10), wetlands (11), rhizosphere (12), and animal guts (13, 14), as free-living organisms or symbionts of eukaryotes. Verrucomicrobia isolates are metabolically diverse, including aerobes, facultative anaerobes, and obligate anaerobes, and they are mostly heterotrophs, using various mono-, oligo-, and poly-saccharides for growth (6, 7, 11, 14-20). Not long ago an autotrophic verrucomicrobial methanotroph (*Methylacidiphilum fumariolicum* SolV) was discovered in acidic thermophilic environments (21).

In marine environments, Verrucomicrobia are also ubiquitous (22) and suggested to have a key role as polysaccharide degraders (23, 24). Genomic insights gained through sequencing single cells (24) or extracting Verrucomicrobia bins from metagenomes (25) have revealed high abundances of glycoside hydrolase genes, providing more evidence for their critical roles in C cycling in marine environments.

In freshwater, Verrucomicrobia have been suggested to degrade glycolate (26) and polysaccharides (24). The abundance of some phylum members was favored by high nutrient availabilities (27, 28), cyanobacterial blooms (29), low pH, high temperature, high hydraulic retention time (30), and more labile DOC (5). To date, there are very few freshwater Verrucomicrobia isolates, including *Verrucomicrobium spinosum* (31) and several *Prosthecobacter* spp. (6). Physiological studies showed that they are aerobes, primarily using carbohydrates, but not amino acids, alcohols, or rarely organic acids for growth. However, these few cultured isolates only represent a single clade within subdivision 1. By contrast, 16S rRNA gene based studies discovered a much wider phylogenic range of freshwater Verrucomicrobia, including subdivisions 1, 2, 3, 4, 5, and 6 (3-5, 24, 32). Due to the very few cultured representatives and few available genomes from this freshwater lineage, the ecological functions of the vast uncultured freshwater Verrucomicrobia are largely unknown.

In this study, we sequenced a total of 184 metagenomes in a time-series study of two lakes with contrasting characteristics, particularly differing in C source, nutrient availabilities, and pH. We recovered a total of 19 Verrucomicrobia draft genomes spanning subdivision 1, 2, 3, and 4 of the seven previously defined Verrucomicrobia subdivisions. We inferred their metabolisms, revealed their adaptation to C and nutrient conditions, and uncovered some interesting and novel features, including a novel putative porin-multiheme cytochrome *c* system that may be involved in extracellular electron transfer. The gained insights advanced our understanding of the ecophysiology, and suggested potential roles in C cycling and ecological niches of this ubiquitous freshwater bacterial group.

## RESULTS AND DISCUSSION

### Comparison of the two lakes

The two studied lakes exhibited contrasting characteristics (Table 1). The most notable difference is the primary C source and nutrient availabilities. Mendota is an urban eutrophic lake with most of its C being autochthonous (in-lake produced through photosynthesis). By contrast, Trout Bog is a nutrient-poor dystrophic lake, surrounded by temperate forests and sphagnum mats, thus receiving large amounts of terrestrially-derived allochthonous C that is rich in humic and fulvic acids. Compared to Mendota, Trout Bog features higher DOC levels, but is more limited in nutrient availability, with much higher DOC:TN and DOC:TP ratios (Table 1). Nutrient limitation in Trout Bog is even more extreme than revealed by these ratios because much of the N and P is tied up in complex dissolved organic matter. In addition, Trout Bog has lower oxygenic photosynthesis due to decreased photosynthetically active radiation (PAR) as a result of absorption by DOC (33). Together with the consumption of dissolved oxygen by heterotrophic respiration, oxygen levels decrease quickly with depth in the water column in Trout Bog. Dissolved oxygen levels are below detection in the hypolimnion nearly year-round (34). Due to these contrasts, we expected to observe differences in bacterial C and nutrient use, as well as differences reflecting the electron acceptor conditions between these two lakes. Hence, the retrieval of numerous Verrucomicrobia draft genomes in the two lakes not only allows the prediction of their general functions in freshwater, but also provides an opportunity to study their ecophysiological adaptation to the local environmental differences.

**TABLE 1.** Lakes included in this study^*a*^

| Lake | Mendota | Trout Bog |
| --- | --- | --- |
| GPS location | 43.100°N, 89.405°W | 46.041°N, 89.686°W |
| Lake type | Drainage lake | Seepage lake |
| Surface area (ha) | 3938 | 1.1 |
| Mean depth (m) | 12.8 | 5.6 |
| Max depth (m) | 25.3 | 7.9 |
| pH | 8.3 | 5.2 |
| Primary carbon source | Phytoplankton | Terrestrial subsidies |
| DOC (mg/L) | 5.0 | 20.0 |
| Total N (mg/L) | 1.5 | 1.3 |
| Total P (µg/L) | 131 | 71 |
| DOC/N | 3.3 | 15.6 |
| DOC/P | 38.0 | 281.9 |
| Trophic state | Eutrophic | Dystrophic |

<sup>a</sup>Data from NTL-LTER https://lter.limnology.wisc.edu, averaged from the study years. DOC = Dissolved organic carbon. N = Nitrogen. P = Phosphorus.

### Verrucomicrobia draft genome retrieval and their distribution patterns

A total of 184 metagenomes were generated from samples collected across multiple years, including 94 from the top 12 m of Mendota (mostly consisting of the epilimnion layer, therefor referred to as “ME”), 45 from Trout Bog epilimnion (“TE”), and 45 from Trout Bog hypolimnion (“TH”). Three combined assemblies were generated by co-assembling reads from all metagenomes within the ME, TE, and TH groups, respectively. Using the binning facilitated by tetranucleotide frequency and relative abundance patterns over time, a total of 19 Verrucomicrobia metagenome-assembled genomes (MAGs) were obtained, including eight from the combined assembly of ME, three from the combined assembly of TE, and eight from the combined assembly of TH (Table 2). The 19 MAGs exhibited a clustering of their tetranucleotide frequency largely based on the two lakes (**Fig. S1**), suggesting distinct overall genomic signatures associated with each system.

**TABLE 2.**
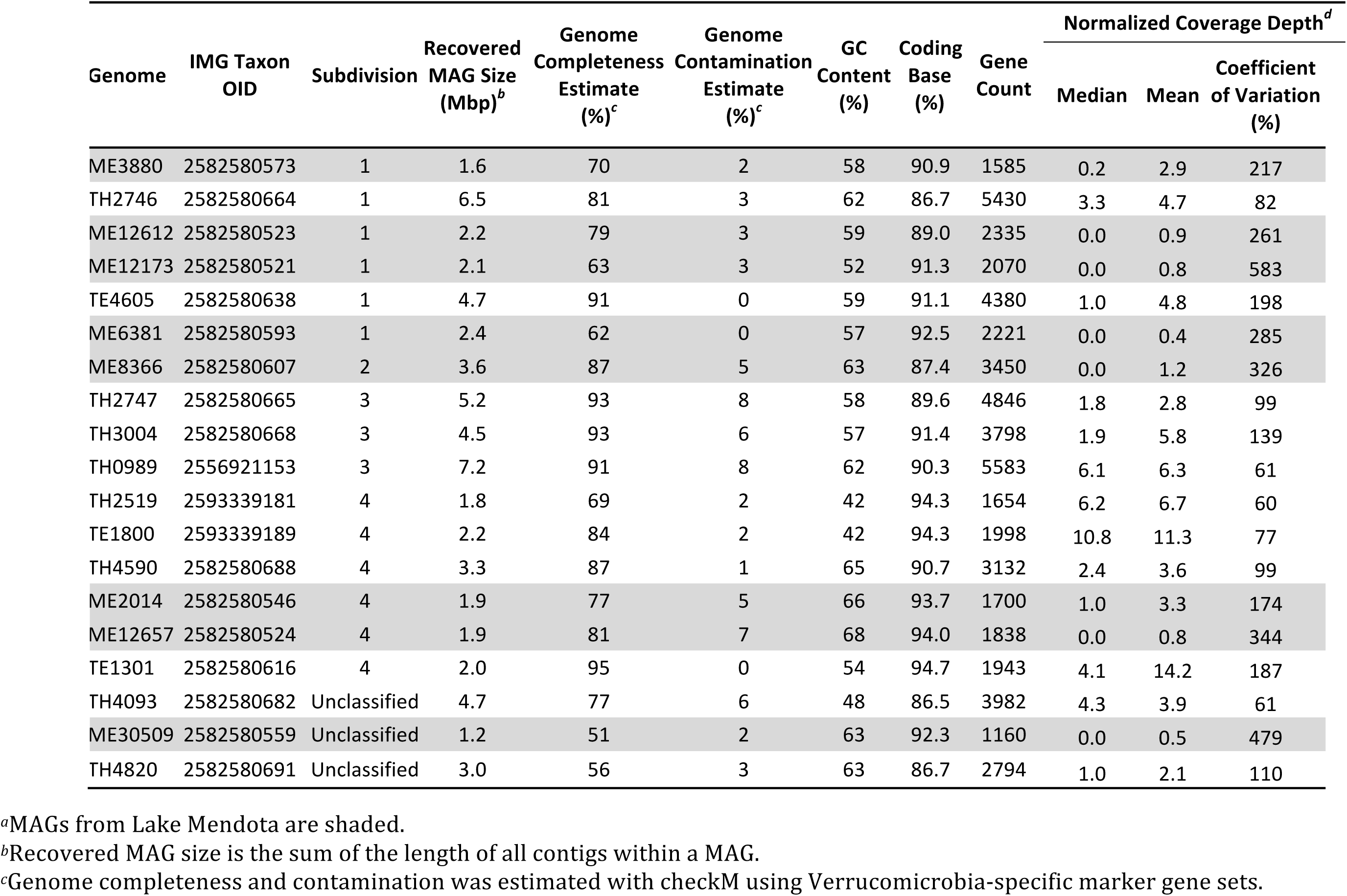
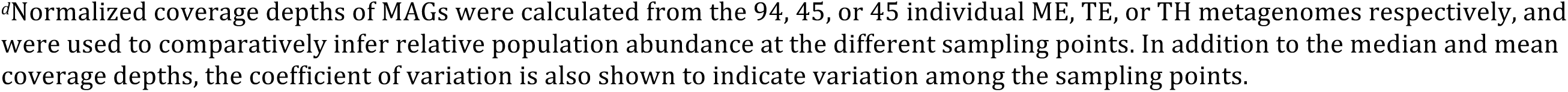
Summary of Verrucomicrobia MAGs^*a*^

| Genome | IMG Taxon<br>OID | Subdivision | Recovered<br>MAG Size<br>(Mbp)<sup>b</sup> | Genome<br>Completeness<br>Estimate<br>(%)<sup>c</sup> | Genome<br>Contamination<br>Estimate<br>(%)<sup>c</sup> | GC<br>Content<br>(%) | Coding<br>Base<br>(%) | Gene<br>Count | Normalized Coverage Depth<sup>d</sup> |  |  |
| --- | --- | --- | --- | --- | --- | --- | --- | --- | --- | --- | --- |
|  |  |  |  |  |  |  |  |  | Median | Mean | Coefficient<br>of Variation<br>(%) |
| ME3880 | 2582580573 | 1 | 1.6 | 70 | 2 | 58 | 90.9 | 1585 | 0.2 | 2.9 | 217 |
| TH2746 | 2582580664 | 1 | 6.5 | 81 | 3 | 62 | 86.7 | 5430 | 3.3 | 4.7 | 82 |
| ME12612 | 2582580523 | 1 | 2.2 | 79 | 3 | 59 | 89.0 | 2335 | 0.0 | 0.9 | 261 |
| ME12173 | 2582580521 | 1 | 2.1 | 63 | 3 | 52 | 91.3 | 2070 | 0.0 | 0.8 | 583 |
| TE4605 | 2582580638 | 1 | 4.7 | 91 | 0 | 59 | 91.1 | 4380 | 1.0 | 4.8 | 198 |
| ME6381 | 2582580593 | 1 | 2.4 | 62 | 0 | 57 | 92.5 | 2221 | 0.0 | 0.4 | 285 |
| ME8366 | 2582580607 | 2 | 3.6 | 87 | 5 | 63 | 87.4 | 3450 | 0.0 | 1.2 | 326 |
| TH2747 | 2582580665 | 3 | 5.2 | 93 | 8 | 58 | 89.6 | 4846 | 1.8 | 2.8 | 99 |
| TH3004 | 2582580668 | 3 | 4.5 | 93 | 6 | 57 | 91.4 | 3798 | 1.9 | 5.8 | 139 |
| TH0989 | 2556921153 | 3 | 7.2 | 91 | 8 | 62 | 90.3 | 5583 | 6.1 | 6.3 | 61 |
| TH2519 | 2593339181 | 4 | 1.8 | 69 | 2 | 42 | 94.3 | 1654 | 6.2 | 6.7 | 60 |
| TE1800 | 2593339189 | 4 | 2.2 | 84 | 2 | 42 | 94.3 | 1998 | 10.8 | 11.3 | 77 |
| TH4590 | 2582580688 | 4 | 3.3 | 87 | 1 | 65 | 90.7 | 3132 | 2.4 | 3.6 | 99 |
| ME2014 | 2582580546 | 4 | 1.9 | 77 | 5 | 66 | 93.7 | 1700 | 1.0 | 3.3 | 174 |
| ME12657 | 2582580524 | 4 | 1.9 | 81 | 7 | 68 | 94.0 | 1838 | 0.0 | 0.8 | 344 |
| TE1301 | 2582580616 | 4 | 2.0 | 95 | 0 | 54 | 94.7 | 1943 | 4.1 | 14.2 | 187 |
| TH4093 | 2582580682 | Unclassified | 4.7 | 77 | 6 | 48 | 86.5 | 3982 | 4.3 | 3.9 | 61 |
| ME30509 | 2582580559 | Unclassified | 1.2 | 51 | 2 | 63 | 92.3 | 1160 | 0.0 | 0.5 | 479 |
| TH4820 | 2582580691 | Unclassified | 3.0 | 56 | 3 | 63 | 86.7 | 2794 | 1.0 | 2.1 | 110 |

<sup>a</sup>MAGs from Lake Mendota are shaded.
<sup>b</sup>Recovered MAG size is the sum of the length of all contigs within a MAG.
<sup>c</sup>Genome completeness and contamination was estimated with checkM using Verrucomicrobia-specific marker gene sets.
<sup>d</sup>Normalized coverage depths of MAGs were calculated from the 94, 45, or 45 individual ME, TE, or TH metagenomes respectively, and were used to comparatively infer relative population abundance at the different sampling points. In addition to the median and mean coverage depths, the coefficient of variation is also shown to indicate variation among the sampling points.

Genome completeness of the 19 MAGs ranged from 51% to 95%, as determined by checkM (35). Phylogenetic analysis of these MAGs using a concatenated alignment of their conserved genes indicates that they span a wide phylogenetic spectrum and distribute in subdivisions 1, 2, 3, and 4 of the seven previously defined Verrucomicrobia subdivisions (5, 21, 36) (Fig. 1), as well as three unclassified Verrucomicrobia MAGs.

**Fig. 1.**
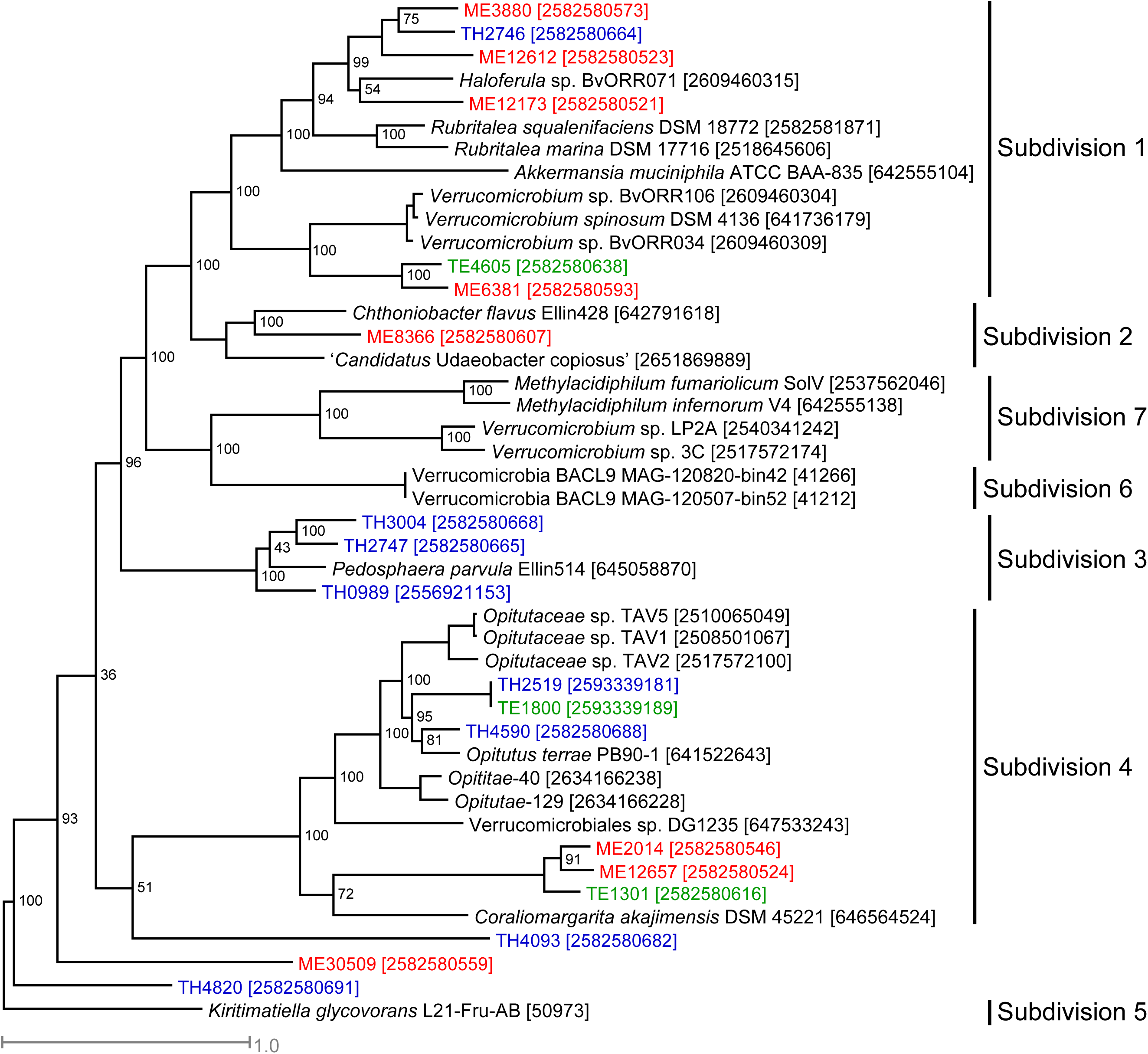
Phylogenetic tree constructed with a concatenated alignment of protein sequences from five conserved essential single-copy genes (represented by TIGR01391, TIGR01011, TIGR00663, TIGR00460, and TIGR00362) that were recovered in all Verrucomicrobia MAGs. ME, TE and TH MAGs are labeled with red, green and blue, respectively. Genome ID in IMG or NCBI is indicated in the bracket. The outgroup is *Kiritimatiella glycovorans* L21-Fru-AB, which was initially assigned to subdivision 5, but this subdivision was recently proposed as a novel sister phylum to Verrucomicrobia (67).

Presently available freshwater Verrucomicrobia isolates are restricted to subdivision 1. The recovered MAGs allow the inference of metabolisms and ecology of a considerable diversity within uncultured freshwater Verrucomicrobia. Notably, all MAGs from subdivision 3 were recovered from TH, and all MAGs from subdivision 1, except TH2746, were from the epilimnion (either ME or TE), indicating differences in phylogenetic distribution between lakes and between layers within a lake.

We used normalized coverage depth of MAGs within individual metagenomes collected at different sampling time points and different lakes/layers to comparatively infer relative population abundance across time and space (see detailed coverage depth estimation in **Supplementary Text**). Briefly, we mapped reads from each metagenome to MAGs with a minimum identity of 95%, and used the number of mapped reads to calculate the relative abundance for each MAG based on coverage depth per contig and several normalization steps. Thus, we assume that each MAG represents a distinct population within the lake-layer from which it was recovered (37, 38). This estimate does not directly indicate the actual relative abundance of these populations within the total community per se; rather it allows us to compare population abundance levels from different lakes and sampling occasions within the set of 19 MAGs. This analysis indicates that Verrucomicrobia populations in Trout Bog were proportionally more abundant and persistent over time compared to those in Mendota in general (Table 2). Verrucomicrobia populations in Mendota boosted their abundances once to a few times during the sampling season and diminished to extremely low levels for the remainder of the sampling season (generally May to November), as reflected by the low median coverage depth of Mendota MAGs and their large coefficient of variation (Table 2)

### Saccharolytic life style and adaptation to different C sources

Verrucomicrobia isolates from different environments are known to grow on various mono-, oligo-, and poly-saccharides, but are unable to grow on amino acids, alcohols, or most organic acids (6, 7, 11, 14-20, 39). Culture-independent research suggests marine Verrucomicrobia as candidate polysaccharide degraders with large number of genes involved in polysaccharide utilization (23-25).

In the 19 Verrucomicrobia MAGs, we observed rich arrays of glycoside hydrolase (GH) genes, representing a total of 78 different GH families acting on diverse polysaccharides (**Fig. S2**). Although these genomes have different degrees of completeness, genome completeness was not correlated with the number of GH genes recovered (correlation coefficient = 0.312, p-value = 0.194), or the number of GH families represented in each MAG (i.e. GH diversity, correlation coefficient = 0.278, p-value = 0.250). To compare GH abundance among MAGs, we normalized GH occurrence frequencies by the total number of genes in each MAG to estimate the percentage of genes annotated as GHs (i.e. GH coding density) to account for the different genome size and completeness. This normalization assumes GH genes are randomly distributed between the recovered and the missing parts of the genome, and it allows us to make some general comparison among these MAGs. GH coding density ranged from 0.4% to 4.9% for these MAGs (Fig. 2a), and in general, was higher in Trout Bog MAGs than in Mendota MAGs. Notably, six TH MAGs had extremely high (∼4%) GH coding densities (Fig. 2a), with each MAG harboring 119-239 GH genes, representing 36-59 different GH families (Fig. 3 **and S2**). Although GH coding density in most ME genomes in subdivisions 1 and 2 was relatively low (0.4-1.6%), it was still higher than in many other bacterial groups (24).

**Fig. 2.**
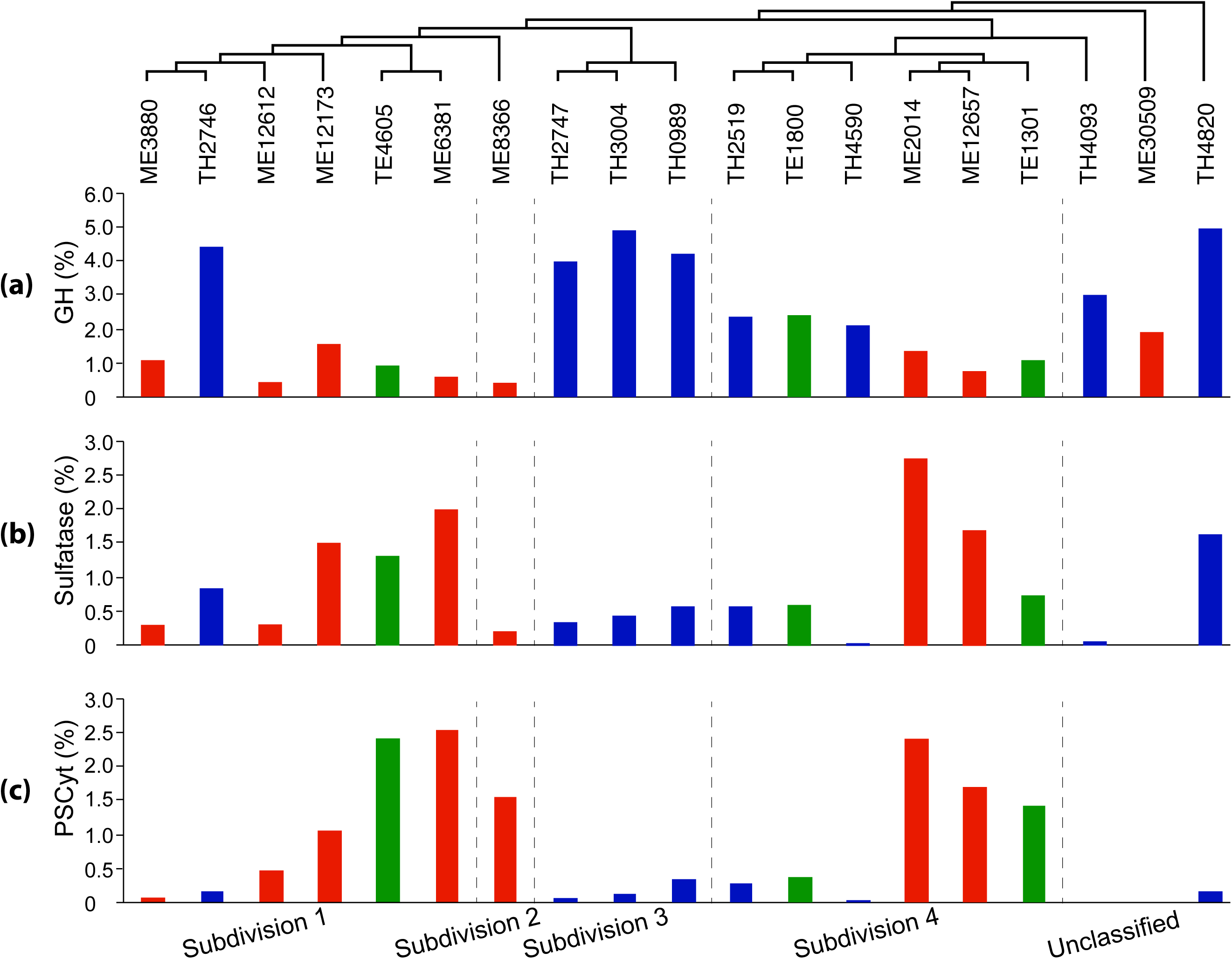
Coding densities of glycoside hydrolase genes (a), sulfatase genes (b) and Planctomycete-specific cytochrome c (PSCyt)-containing genes (c). Data from ME, TE and TH MAGs are labeled with red, green and blue, respectively. The three plots share the same x-axis label indicated by the genome clustering on the top, which is is based on a subtree extracted from the phylogenetic tree in Fig. 1 to indicate the phylogenetic relatedness of the 19 MAGs. The vertical dashed lines divide these MAGs to different subdivisions.

The GH abundance and diversity within a genome may determine the width of the substrate spectrum and/or the complexity of carbohydrates used by that organism. For example, there are 20 GH genes in the *Rubritalea marina* genome, and this marine verrucomicrobial aerobe only uses a limited spectrum of carbohydrate monomers and dimers, but not the majority of (poly)saccharides tested (15). By contrast, 164 GH genes are present in the *Opitutus terrae* genome, and this soil verrucomicrobial anaerobe can thus grow on a wider range of mono-, di- and poly-saccharides (16). Therefore, it is plausible that the GH-rich Trout Bog Verrucomicrobia populations may be able to use a wider range of more complex polysaccharides than the Mendota populations.

The 10 most abundant GH families in these Verrucomicrobia MAGs include GH2, 29, 78, 95, and 106 (Fig. 3). These specific GHs were absent or at very low abundances in marine Verrucomicrobia genomes (24, 25), suggesting a general difference in carbohydrate substrate use between freshwater- and marine Verrucomicrobia. Hierarchical clustering of MAGs based on overall GH abundance profiles indicated a grouping pattern largely separated by lake (**Fig. S3**). Prominently over-represented GHs in most Trout Bog MAGs include GH2, GH29, 78, 95, and 106. By contrast, over-represented GHs in the Mendota MAGs are GH13, 20, 33, 57, and 77, which have different substrate spectra from GHs over-represented in the Trout Bog MAGs. Therefore, the patterns in GH functional profiles may suggest varied carbohydrate substrate preferences and ecological niches occupied by Verrucomicrobia, probably reflecting the different carbohydrate composition derived from different sources between Mendota and Trout Bog.

Overall, GH diversity and abundance profile may reflect the DOC availability, chemical variety and complexity, and suggest microbial adaptation to different C sources in the two ecosystems. We speculate that the rich arrays of GH genes, and presumably broader substrate spectra of Trout Bog populations, partly contribute to their higher abundance and persistence over the sampling season (Table 2), as they are less likely impacted by fluctuations of individual carbohydrates. By contrast, Mendota populations with fewer GHs and presumably more specific substrate spectra are relying on autochthonous C and therefore exhibit a “bloom-and-bust” abundance pattern (Table 2) that might be associated with cyanobacterial blooms as previous suggested (29). On the other hand, bogs experience seasonal phytoplankton blooms (40, 41) that introduce brief pulses of autochthonous C to these otherwise allochthonous-driven systems. Clearly, much remains to be learned about the routes through which C is metabolized by bacteria in such lakes, and comparative genomics is a novel way to use the organisms to tell us about C flow through the ecosystem.

### Other genome features of the saccharide-degrading life style

Seven Verrucomicrobia MAGs spanning subdivisions 1, 2, 3, and 4 possess genes needed to construct bacterial microcompartments (BMCs), which are quite rare among studied bacterial lineages. Such BMC genes in Planctomycetes are involved in the degradation of plant and algal cell wall sugars, and are required for growth on L-fucose, L-rhamnose and fucoidans (42). Genes involved in L-fucose and L-rhamnose degradation cluster with BMC shell protein-coding genes in the seven Verrucomicrobia MAGs (Fig. 4). This is consistent with the high abundance of *α*-L-fucosidase or *α*-L-rhamnosidase GH genes (represented by GH29, 78, 95, 106) in most of these MAGs (Fig. 3), suggesting the importance of fucose- and rhamnose-containing polysaccharides for these Verrucomicrobia populations.

**Fig. 3.**
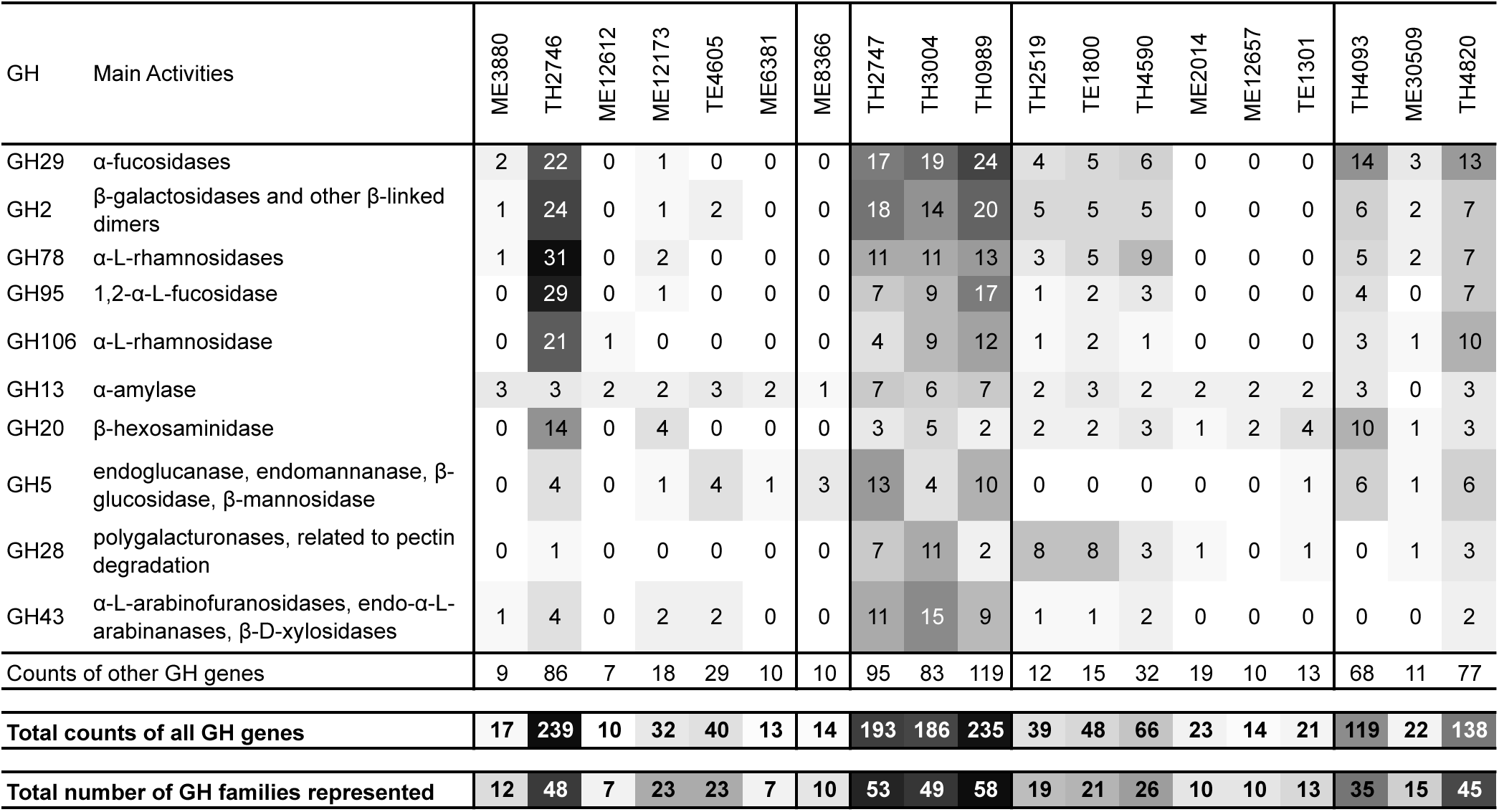
Gene counts for the top 10 most abundant GH families, total gene counts for all GH families, and the number of GH families represented by these genes. MAGs are ordered as in the clustering in Fig. 2.

**Fig. 4.**
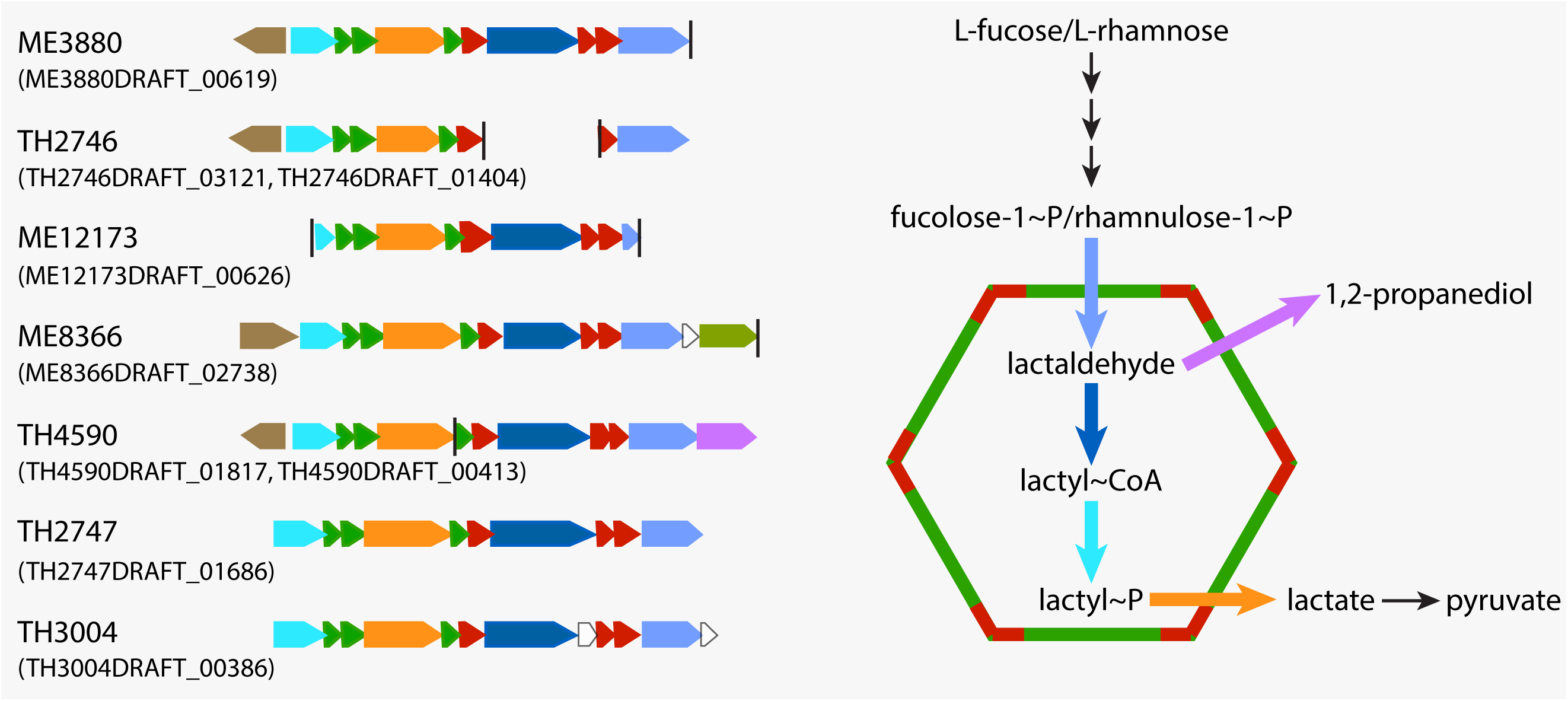
Gene clusters encoding bacterial microcompartments (BMCs) involved in L-fucose and L-rhamnose degradation. The vertical line indicates the end of a contig, and IMG gene locus tag for the first gene in each presented gene cluster is indicated in the parenthesis. The BMC is schematically represented by a hexagon with the two building blocks labeled in red and green, respectively. The two building blocks and reactions inside the BMC are colored according to their encoding genes’ color labels on the left side.

TonB-dependent receptor (TBDR) genes were found in Verrucomicrobia MAGs, and are present at over 20 copies in TE1800 and TH2519. TBDRs are located on the outer cellular membrane of Gram-negative bacteria, usually mediating the transport of iron siderophore complex and vitamin B_12_ across the outer membrane through an active process. More recently, TBDRs were suggested to be involved in carbohydrate transport across the outer membrane by some bacteria that consume complex carbohydrates, and in their carbohydrate utilization (CUT) loci, TBDR genes usually cluster with genes encoding inner membrane transporters, GHs and regulators for efficient carbohydrate transportation and utilization (43). Such novel CUT loci are present in TE1800 and TH2519, with TBDR genes clustering with genes encoding inner membrane sugar transporters, monosaccharide utilization enzymes, and GHs involved in the degradation of pectin, xylan, and fucose-containing polymers (Fig. 5). Notably, most GHs in the CUT loci are predicted to be extracellular or outer membrane proteins (Fig. 5), catalyzing extracellular hydrolysis reactions to release mono- and oligo-saccharides, which are transported across the outer membrane by TBDR proteins. Therefore, such CUT loci may allow these verrucomicrobial populations to coordinately and effectively scavenge the hydrolysis products before they diffuse away.

**Fig. 5.**
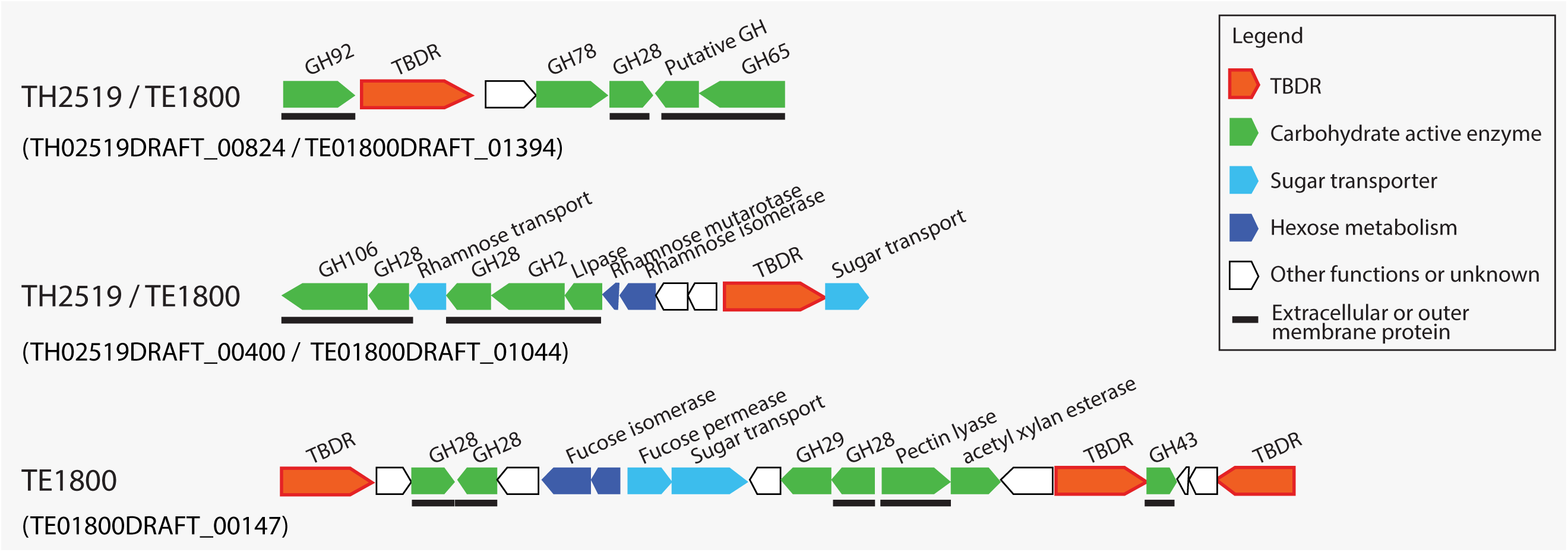
Gene clusters encoding putative tonB-dependent carbohydrate utilization (CUT) loci. IMG gene locus tag for the first gene in each presented gene cluster is indicated in the parenthesis. The horizontal solid lines below genes indicate predicted extracellular or outer membrane proteins.

Genes encoding for inner membrane carbohydrate transporters are abundant in Verrucomicrobia MAGs (**Fig. S4**). The Embden-Meyerhof pathway for glucose degradation, as well as pathways for degrading a variety of other sugar monomers, including galactose, rhamnose, fucose, xylose, and mannose, were recovered (complete or partly-complete) in most MAGs (Fig. 6). As these sugars are abundant carbohydrate monomers in plankton and plant cell walls, the presence of these pathways together with GH genes suggest that these Verrucomicrobia populations may use plankton- and plant-derived saccharides. Machinery for pyruvate degradation to acetyl-CoA and the TCA cycle are also present in most MAGs. These results are largely consistent with their hypothesized role in carbohydrate degradation and previous studies on Verrucomicrobia isolates.

**Fig. 6.**
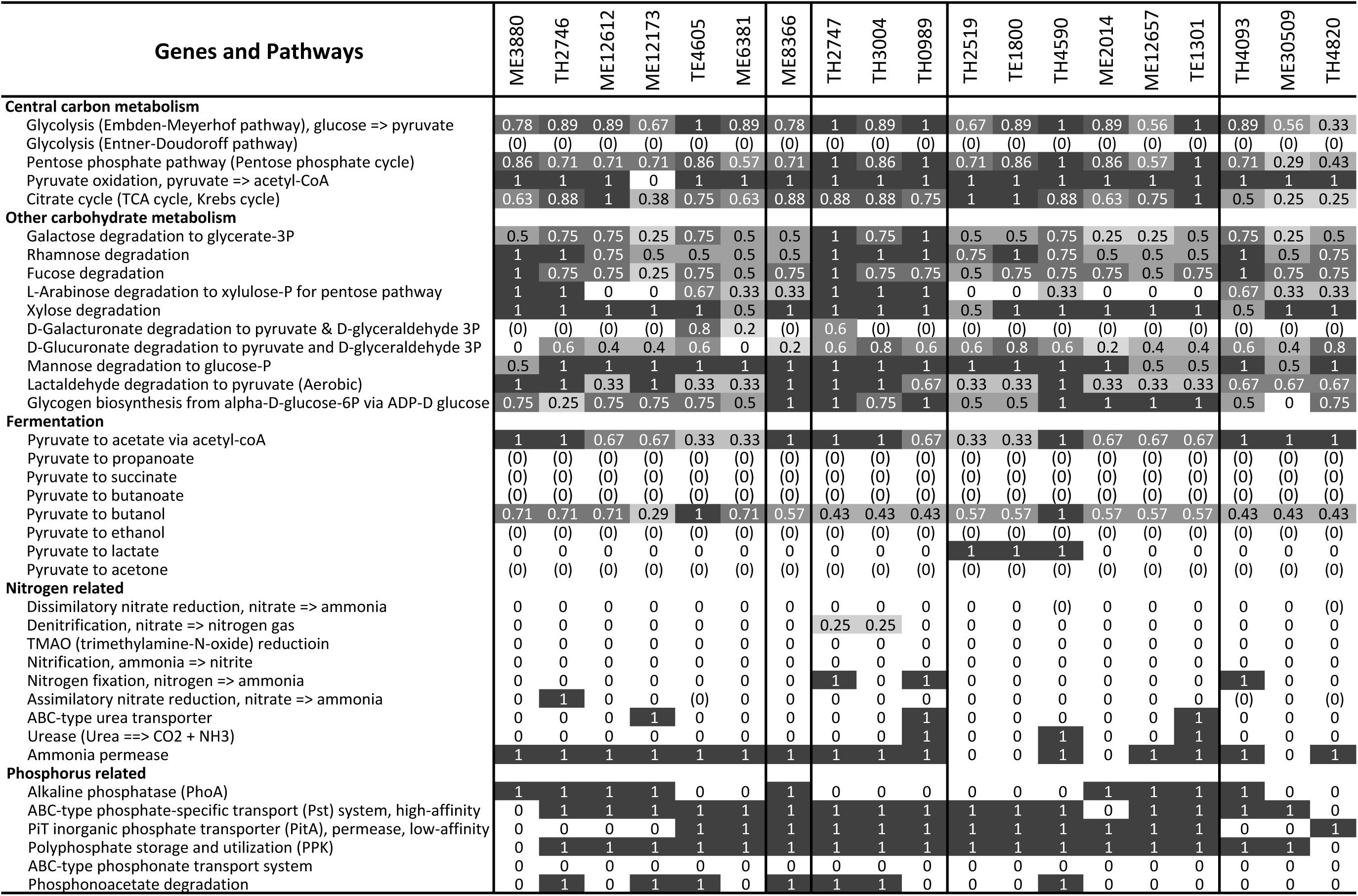
Completeness estimates of key metabolic pathways. MAGs are ordered as in the clustering in Fig. 2. Completeness value of “1” indicates a pathway is complete; “0” indicates no genes were found in that pathway; and “(0)” indicates that although some genes in a pathway are present, the pathway is likely absent because signature genes for that pathway were not found in that draft genome AND signature genes are missing in more than two thirds of all draft genomes.

Notably, a large number of genes encoding proteins belonging to a sulfatase family (pfam00884) are present in the majority of MAGs (Fig. 2b), similar to the high representation of these genes in marine Verrucomicrobia genomes (24, 25). Sulfatases hydrolyze sulfate esters, which are rich in sulfated polysaccharides. In general, sulfated polysaccharides are abundant in marine algae and plants (mainly in seaweeds) (44), but have also been found in some freshwater cyanobacteria (45) and plant species (46). Sulfatase genes in our Verrucomicrobia MAGs were often located in the same neighborhood as genes encoding for extracellular proteins with a putative pectin lyase activity, proteins with a carbohydrate-binding module (pfam13385), GHs, and proteins with PSCyt domains (Fig. 2c and discussed later). Their genome context lends support for the participation of these genes in C and sulfur cycling by degrading sulfated polysaccharides, which can serve as an abundant source of sulfur for cell biosynthesis as well as C for energy and growth.

Previously, freshwater Verrucomicrobia were suggested to use the algal exudate glycolate in humic lakes, based on the retrieval of genes encoding subunit D (*glcD*) of glycolate oxidase, which converts glycolate to glyoxylate (26). However, these recovered genes might not be bona fide *glcD* due to the lack of other essential subunits as revealed in our study (see **Supplementary Text**). Among the MAGs, only TE4605 possesses all three essential subunits of glycolate oxidase (*glcDEF)* (**Fig. S5**). However, genetic context analysis suggests that TE4605 likely uses glycolate for amino acid assimilation, instead of energy generation (**Fig. S5** and **Supplementary Text**). These results are consistent with the absence of the glyoxylate shunt and especially the malate synthase, which converts glyoxylate to malate to be used through the TCA cycle for energy generation in the 19 Verrucomicrobia MAGs (**Fig. S6**). Therefore, Verrucomicrobia populations represented by the 19 MAGs are not likely key players in glycolate degradation, but more likely important (poly)saccharide-degraders in freshwater, as suggested by the high abundance of GH, sulfatase, and carbohydrate transporter genes, metabolic pathways for degrading diverse carbohydrate monomers, and other genome features adapted to the saccharolytic life style.

### Nitrogen (N) metabolism and adaptation to different N availabilities

Most Verrucomicrobia MAGs in our study do not appear to reduce nitrate or other nitrogenous compounds, and they seem to uptake and use ammonia (Fig. 6), and occasionally amino acids (**Fig. S4**), as an N-source. Further, some Trout Bog populations may have additional avenues to generate ammonia, including genetic machineries for assimilatory nitrate reduction in TH2746, nitrogenase genes for nitrogen fixation and urease genes in some of the Trout Bog MAGs (Fig. 6), probably as adaptions to N-limited conditions in Trout Bog.

Although Mendota is a eutrophic lake, N can become temporarily limiting during the high-biomass period when N is consumed by large amounts of phytoplankton and bacterioplankton (47). For some bacteria, when N is temporarily limited while C is in excess, cells convert and store the extra C as biopolymers. For example, the verrucomicrobial methanotroph *M. fumariolicum* SolV accumulated a large amount of glycogen (up to 36% of the total dry weight of cells) when the culture was N-limited (48). Similar to this verrucomicrobial methanotroph, genes in glycogen biosynthesis are present in most MAGs from Mendota and Trout Bog (Fig. 6). Indeed, a glycogen synthesis pathway is also present in most genomes of cultivated Verrucomicrobia in the public database (data not shown), suggesting that glycogen accumulation might be a common feature for this phylum to cope with the changing pools of C and N in the environment and facilitate their survival when either is temporally limited.

### Phosphorus (P) metabolism and other metabolic features

Verrucomicrobia populations represented by these MAGs may be able to survive under low P conditions, as suggested by the presence of genes responding to P limitation, such as the two-component regulator (*phoRB*), alkaline phosphatase (*phoA*), phosphonoacetate hydrolase (*phnA*), and high-affinity phosphate-specific transporter system (*pstABC*) (Fig. 6). Detailed discussion in P acquisition and metabolism and other metabolic aspects, such as acetate metabolism, sulfur metabolism, oxygen tolerance, and the presence of the alternative complex III and cytochrome *c* oxidase genes in the oxidative phosphorylation pathway, are discussed in the Supplementary Text (**Fig. S6**).

### Anaerobic respiration and a putative porin-multiheme cytochrome *c* system

Respiration using alternative electron acceptors is important for overall lake metabolism in the DOC-rich humic Trout Bog, as the oxygen levels decrease quickly with depth in the water column. We therefore searched for genes involved in anaerobic respiration, and found that genes in the dissimilatory reduction of nitrate, nitrite, sulfate, sulfite, DMSO, and TMAO are largely absent in all MAGs (**Supplementary Text, Fig. S6**). Compared to those anaerobic processes, genes for dissimilatory metal reduction are less well understood. In more extensively studied cultured iron [Fe(III)] reducers, outer surface *c*-type cytochromes (cyt*c*), such as OmcE and OmcS in *Geobacter sulfurreducens* are involved in Fe(III) reduction at the cell outer surface (49). Further, a periplasmic multiheme cytochrome *c* (MHC, e.g. MtrA in *Shewanella oneidensis* and OmaB/OmaC in *G. sulfurreducens*) can be embedded into a porin (e.g. MtrB in *S. oneidensis* and OmbB/OmbC in *G. sulfurreducens*), forming a porin-MHC complex as an extracellular electron transfer (EET) conduit to reduce extracellular Fe(III) (50, 51). Such outer surface cyt*c* and porin-MHC systems involved in Fe(III) reduction were also suggested to be important in reducing the quinone groups in humic substances (HS) at the cell surface (52-54). The reduced HS can be re-oxidized by Fe(III) or oxygen, thus HS can serve as electron shuttles to facilitate Fe(III) reduction (55, 56) or as regenerable electron acceptors at the anoxic-oxic interface or over redox cycles (57).

Outer surface cyt*c* or porin-MHC systems homologous to the ones in *G. sulfurreducens* and *S. oneidensis* are not present in Verrucomicrobia MAGs. Instead, we identified a novel porin-coding gene clustering with MHC genes in six MAGs (Fig. 7). These porins were predicted to have at least 20 transmembrane motifs, and their adjacent cyt*c* were predicted to be periplasmic proteins with eight conserved heme-binding sites. In several cases, a gene encoding an extracellular MHC is also located in the same gene cluster. As their gene organization is analogous to the porin-MHC gene clusters in *G. sulfurreducens* and *S. oneidensis*, we hypothesize that these genes in Verrucomicrobia may encode a novel porin-MHC complex involved in EET.

**Fig. 7.**
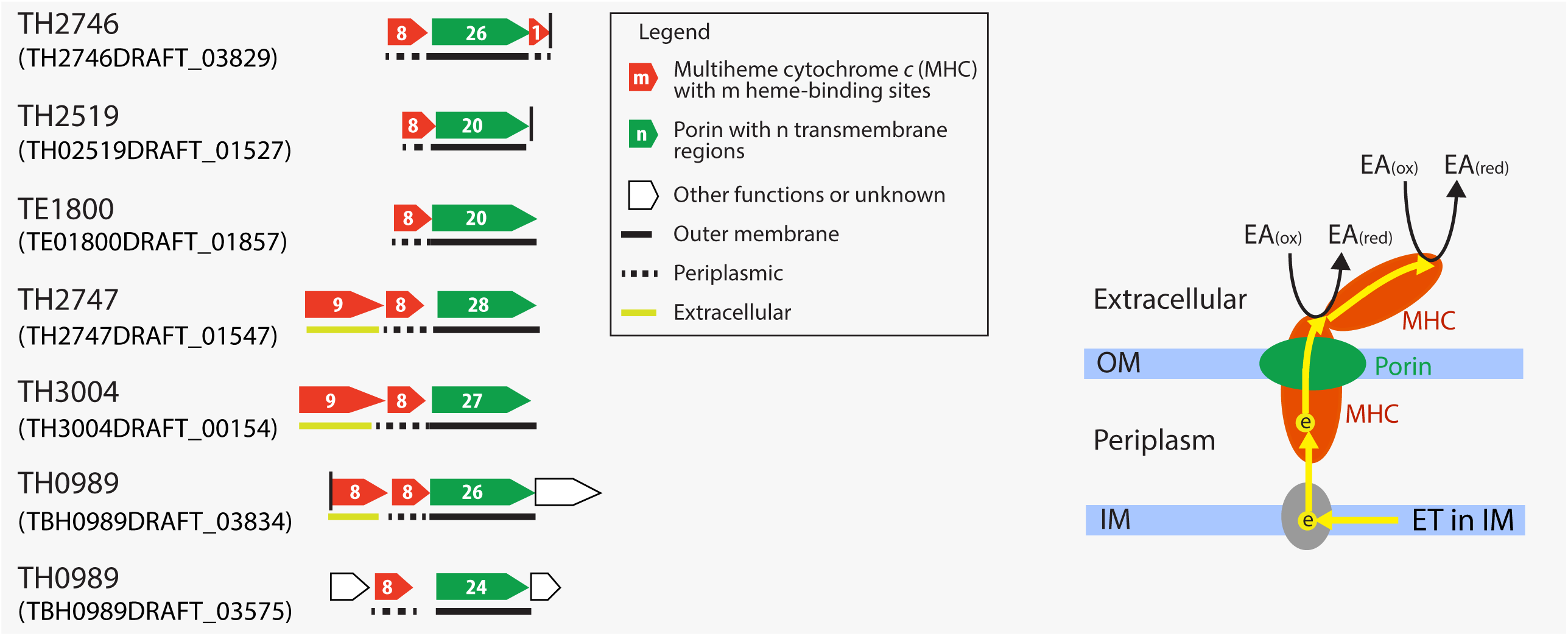
Gene clusters encoding putative porin-multiheme cytochrome c complex (PCC). IMG gene locus tag for the first gene in each presented gene cluster is indicated in the parenthesis. The vertical line indicates the end of a contig, and horizontal lines below genes indicate predicted cellular locations of their encoded proteins. These putative PCC genes are in 18.1, 9.0, 6.1, 18.4, 70.0, 10.6 and 10.8 kbp long contigs, respectively. A hypothesized model of extracellular electron transfer is shown on the right with yellow arrows indicating electron flows. “IM” and “OM” refer to inner and outer membranes, respectively, “ET in IM” refers to electron transfer in the inner membrane, and “EA_(ox)_” and “EA_(red)_” refer to oxidized and reduced forms of the electron acceptor, respectively.

As these porin-MHC gene clusters are novel, we further confirmed that they are indeed from Verrucomicrobia. Their containing contigs were indeed classified to Verrucomicrobia based on the consensus of the best BLASTP hits for genes on these contigs. Notably, the porin-MHC gene cluster was only observed in MAGs recovered from the HS-rich Trout Bog, especially from the anoxic hypolimnion environment. Searching the NCBI and IMG databases for the porin-MHC gene clusters homologous to those in Trout Bog, we identified homologs in genomes within the Verrucomicrobia phylum, including *Opitutus terrae* PB90-1 isolated from rice paddy soil, *Opitutus* sp. GAS368 isolated from forest soil, “*Candidatus* Udaeobacter copiosus” recovered from prairie soil, Opititae-40 and Opititae-129 recovered from freshwater sediment, and Verrucomicrobia bacterium IMCC26134 recovered from freshwater; some of their residing environments are also rich in HS. Therefore, based on the occurrence pattern of porin-MHC among Verrucomicrobia genomes, we hypothesize that such porin-MHCs might participate in EET to HS in anoxic HS-rich environments, and HS may further shuttle electrons to poorly soluble metal oxides or be regenerated at the anoxic-oxic interface, thereby diverting more C flux to respiration instead of fermentation and methanogenesis, which could impact the overall energy metabolism and green-house gas emission in the bog environment.

### Occurrence of Planctomycete-specific cytochrome *c* and domains

One of the interesting features of Verrucomicrobia and its sister phyla in the PVC superphylum is the presence of a number of novel protein domains in some of their member genomes (58, 59). These domains were initially identified in marine planctomycete *Rhodopirellula baltica* (58) and therefore, were referred to as “Planctomycete-specific”, although some of them were later identified in other PVC members (59). In our Verrucomicrobia MAGs, most genes containing Planctomycete-specific cytochrome *c* domains (PSCyt1 to PSCyt3) also contain other Planctomycete-specific domains (PSD1 through PSD5) with various combinations and arrangements (Fig. 8 and **S7a**). Further, PSCyt2-containing and PSCyt3-containing genes are usually next to two different families of unknown genes, respectively (**Fig. S7b**). Such conserved domain architectures and gene organizations, as well as their high occurrence frequencies in some of the Verrucomicrobia MAGs are intriguing, yet nothing is known about their functions. However, some of the PSCyt-containing genes also contain protein domains identifiable as carbohydrate-binding modules (CBMs), suggesting a role in carbohydrate metabolism (see detailed discussion in **Supplementary Text**).

**Fig. 8.**
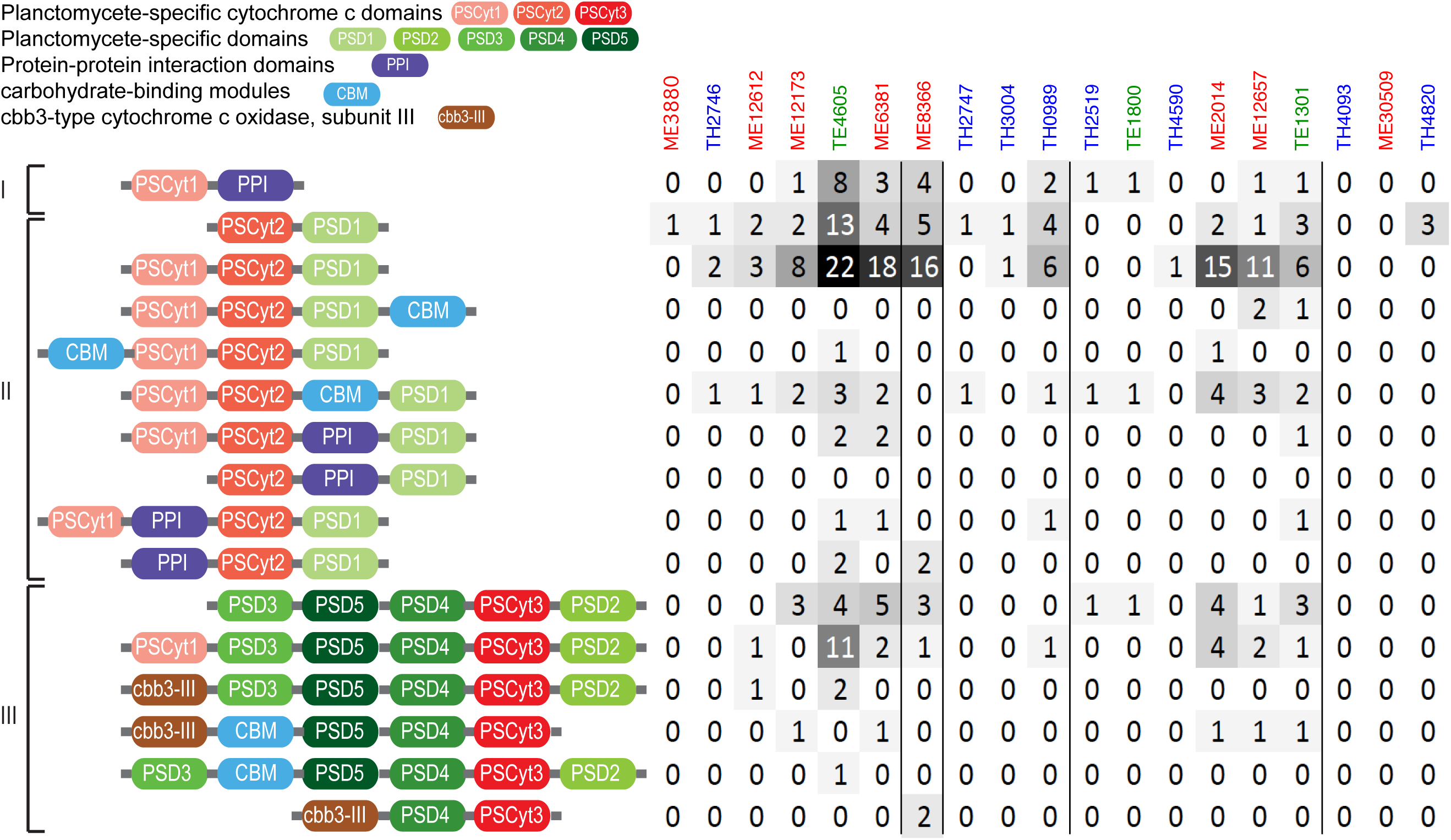
Domain architecture and occurrence of PSCyt-containing genes. Based on the combination of specific PSCyt and PSD domains, these domain structures can be classified into three groups (I, II, and III). “CBM” refers to carbohydrate-binding modules, which include pfam13385 (Laminin_G_3), pfam08531 (Bac_rhamnosid_N), pfam08305 (NPCBM), pfam03422 (CBM_6), and pfam07691 (PA14). “PPI” refers to protein-protein interaction domains, which include pfam02368 (Big_2), pfam00400 (WD40), and pfam00754 (F5_F8_type_C).

The coding density of PSCyt-containing genes indicates that they tend to be more abundant in the epilimnion (either ME or TE) genomes (Fig. 2c) and exhibit an inverse correlation with the GH coding density (*r* = −0.62). Interestingly, sulfatase-coding genes are often in the neighborhood of PSCyt-containing genes in ME and TE genomes, whereas sulfatase-coding genes often neighbor with GH genes in TH genomes. The genomic context suggests PSCyt-containing gene functions somewhat mirror those of GHs (although their reaction mechanisms likely differ fundamentally). However, these PSCyt-containing genes were predicted to be periplasmic or cytoplasmic proteins rather than extracellular or outer membrane proteins. Hence, if they are indeed involved in carbohydrate degradation, they likely act on mono- or oligomers that can be transported into the cell. Further, the distribution patterns of GH versus PSCyt-containing genes between the epilimnion and hypolimnion may reflect the difference in oxygen availability and their carbohydrate substrate complexity between the two layers, suggesting some niche differentiation within Verrucomicrobia in freshwater systems. Therefore, we suggest that a combination of carbohydrate composition, electron acceptor availability and C accessibility drive gene distributions in these populations.

### Summary

Verrucomicrobia MAGs recovered from the two contrasting lakes greatly expanded the known genomic diversity of freshwater Verrucomicrobia, revealed the ecophysiology and some interesting adaptive features of this ubiquitous yet less understood freshwater lineage. The overrepresentation of GH, sulfatase, and carbohydrate transporter genes, the genetic potential to use various sugars, and the microcompartments for fucose and rhamnose degradation suggest that they are potentially (poly)saccharide degraders in freshwater. Most of the MAGs encode machineries to cope with the changing availability of N and P and can survive nutrient limitation. Despite these generalities, these Verrucomicrobia differ significantly between lakes in the abundance and functional profiles of their GH genes, which may reflect different C sources of the two lakes. Interestingly, a number of MAGs in Trout Bog possess gene clusters potentially encoding a novel porin-multiheme cytochrome *c* complex, and might be involved in extracellular electron transfer in the anoxic humic-rich environment. Intriguingly, large numbers of Planctomycete-specific cytochrome *c*-containing genes are present in MAGs from the epilimnion, exhibiting nearly opposite distribution patterns with GH genes. Future studies are needed to elucidate the functions of these novel and fascinating genomic features.

In this study, we focused on using genome information to infer ecophysiology of Verrucomicrobia. The rich time-series metagenome dataset and the many diverse microbial genomes recovered in these two lakes also provide an opportunity for the future study of Verrucomicrobia population dynamics in the context of the total community and their interactions with environmental variables and other microbial groups.

As some of the MAGs analyzed here represent first genome representatives of several Verrucomicrobia subdivisions from freshwater, an interesting question is whether populations represented by the MAGs are native aquatic residents and active in aquatic environment, or merely present after having been washed into the lake from surrounding soil. Previous studies on freshwater Verrucomicrobia were largely based on 16S rRNA genes, yet 16S rRNA genes were not recovered in most MAGs, making it difficult to directly link our MAGs to previously identified freshwater Verrucomicrobia. Notably, our MAGs were only distantly related to the ubiquitous and abundant soil Verrucomicrobia, “*Candidatus* Udaeobacter copiosus” (10) (Fig. 1). In addition, Verrucomicrobia were abundant in Trout Bog and other bogs from a five-year bog lake bacterial community composition and dynamics study (60), with average relative abundance of 7.1% and 8.6%, and maximal relative abundance of 25.4% and 39.5% in Trout Bog epilimnion and hypolimnion respectively. Since the MAGs were presumably from the most abundant Verrucomicrobia populations, they were not likely soil immigrants due to their high abundance in the aquatic environment. To confirm their aquatic origin, future experiments should be designed to test their activities and physiology in the aquatic environment based on the genomic insights gained in this study.

## MATERIALS AND METHODS

### Study sites

Samples for metagenome sequencing were collected from two temperate lakes in Wisconsin, USA, Lake Mendota and Trout Bog Lake, during ice-off periods of each year (May to November). Mendota is an urban eutrophic lake with most of its C being autochthonous (in-lake produced), whereas Trout Bog is a small, acidic and nutrient-poor dystrophic lake with mostly terrestrially-derived (allochthonous) C. General lake characteristics are summarized in Table 1.

### Sampling

For Mendota, we collected depth-integrated water samples from the surface 12 m (mostly consisting of the epilimnion layer) at 94 time points from 2008 to 2012, and samples were referred to as “ME” (38). For Trout Bog, we collected the integrated hypolimnion layer at 45 time points from 2007 to 2009 and the integrated epilimnion layer at 45 time points from 2007 to 2009, and samples were referred to as “TH” and “TE”, respectively (37). All samples were filtered through 0.22 *µ*m polyethersulfone filters and stored at −80°C until extraction. DNA was extracted from the filters using the FastDNA kit (MP Biomedicals) according to manufacturer’s instruction with some minor modifications as described previously (34).

### Metagenome sequencing, assembly, and draft genome recovery

Details of metagenome sequencing, assembly, and binning were described in Bendall *et al.* (37) and Hamilton et al (61). Briefly, shotgun Illumina HiSeq 2500 metagenome libraries were constructed for each of the DNA samples. Three combined assemblies were generated by co-assembling reads from all metagenomes within the ME, TE, and TH groups, respectively. Binning was conducted on the three combined assemblies to recover “metagenome-assembled genomes” (MAGs) based on the combination of contig tetranucleotide frequency and differential coverage patterns across time points using MetaBAT (62). Subsequent manual curation of MAGs was conducted to remove contigs that did not correlate well with the median temporal abundance pattern of all contigs within a MAG, as described in Bendall *et al.* (37).

### Genome annotation and completeness estimation

MAGs were submitted to the DOE Joint Genome Institute’s Integrated Microbial Genome (IMG) database for gene prediction and function annotation (63). The IMG Taxon Object IDs for Verrucomicrobia MAGs are listed in Table 2. The completeness and contamination of each MAG was estimated using checkM with both the lineage-specific and Verrucomicrobia-specific workflows (35). The Verrucomicrobia-specific workflow provided more accurate estimates (i.e. higher genome completeness and lower contamination) than the lineage-specific workflow when tested on 11 complete genomes of Verrucomicrobia isolates available at IMG during our method validation. We therefore only reported the estimates from Verrucomicrobia-specific workflow (Table 2). MAGs with an estimated completeness lower than 50% were not included in this study.

### Taxonomic and phylogenetic analysis

A total of 19 MAGs were classified to the Verrucomicrobia phylum based on taxonomic assignment by PhyloSift using 37 conserved phylogenetic marker genes (64), as described in Bendall *et al.* (37). A phylogenetic tree was reconstructed from the 19 Verrucomicrobia MAGs and 24 reference genomes using an alignment concatenated from individual protein alignments of five conserved essential single-copy genes (represented by TIGR01391, TIGR01011, TIGR00663, TIGR00460, and TIGR00362) that were recovered in all Verrucomicrobia MAGs. Individual alignments were first generated with MUSCLE (65), concatenated, and trimmed to exclude columns that contain gaps for more than 30% of all sequences. A maximum likelihood phylogenetic tree was constructed using PhyML 3.0 (66), with the LG substitution model and the gamma distribution parameter estimated by PhyML. Bootstrap values were calculated based on 100 replicates. *Kiritimatiella glycovorans* L21-Fru-AB was used as an outgroup in the phylogenetic tree. This bacterium was initially designated as the first (and so far the only) cultured representative of Verrucomicrobia subdivision 5. However, this subdivision was later proposed as a novel sister phylum associated with Verrucomicrobia (67), making it an ideal outgroup for this analysis.

### Estimate of metabolic potential

IMG provides functional annotation based on KO (KEGG orthology) term, COG (cluster of orthologous group), pfam, and TIGRfam. To estimate metabolic potential, we primarily used KO terms due to their direct link to KEGG pathways. COG, pfam, and TIGRfam were also used when KO terms were not available for a function. Pathways are primarily reconstructed according to KEGG modules, and MetaCyc pathway is used if a KEGG module is not available for a pathway. As these MAGs are incomplete genomes, a fraction of genes in a pathway may be missing due to genome incompleteness. Therefore, we estimated the completeness of a pathway as the fraction of recovered enzymes in that pathway (e.g. a pathway is 100% complete if all enzymes in that pathway are encoded by genes recovered in a MAG). As some genes are shared by multiple pathways, signature genes specific for a pathway were used to indicate the presence of a pathway. If signature genes for a pathway were missing in all MAGs, that pathway was likely absent in all genomes. Based on this, we established criteria for estimating pathway completeness in each MAG. If a signature gene in a pathway was present, we report the percentage of genes in the pathway that we found. If a signature gene was absent in a MAG, but present in at least one third of all MAGs (i.e. >=7), we still report the pathway completeness for that MAG in order to account for genome incompleteness. Otherwise, we considered the pathway to be absent (i.e. completeness is 0%).

### Glycoside hydrolase identification

Glycoside hydrolase (GH) genes were identified using the dbCAN annotation tool (http://csbl.bmb.uga.edu/dbCAN/annotate.php) (68) using HMMER search against hidden Markov models (HMMs) built for all GHs, with an E-value cutoff of 1e-7, except GH109, for which we found that the HMM used by dbCAN is pfam01408, which is a small domain at the N-terminus of GH109 proteins, but is not specific for GH109. Therefore, to identify verrucomicrobial GH109, BLASTP was performed using the two GH109 sequences (GenBank accession ACD03864 and ACD04752) from verrucomicrobial *Akkermansia muciniphila* ATCC BAA-835 listed in the CAZy database (http://www.cazy.org), with E-value cutoff of 1e-6 and query sequence coverage cutoff of 50%.

### Other bioinformatic analyses

Protein cellular location was predicted using CELLO v.2.5 (http://cello.life.nctu.edu.tw) (69) and PSORTb v.3.0 (http://www.psort.org/psortb) (70). The beta-barrel structure of outer membrane proteins was predicted by PRED-TMBB (http://bioinformatics.biol.uoa.gr//PRED-TMBB) (71).

## ACKNOWLEDGEMENTS

We thank the North Temperate Lakes Microbial Observatory 2007-2012 field crews, UW-Trout Lake Station, the UW Center for Limnology, and the Global Lakes Ecological Observatory Network for field and logistical support. We give special thanks to past McMahon lab graduate students Ashley Shade, Ryan Newton, Emily Read, and Lucas Beversdorf. We acknowledge efforts by many McMahon Lab undergrads and technicians related to sample collection and DNA extraction, particularly Georgia Wolfe. We personally thank the individual program directors and leadership at the National Science Foundation for their commitment to continued support of long term ecological research.

The project was supported by funding from the United States National Science Foundation Microbial Observatories program (MCB-0702395), the Long Term Ecological Research program (NTL-LTER DEB-1440297) and an INSPIRE award (DEB-1344254). This material is also based upon work that supported by the National Institute of Food and Agriculture, U.S. Department of Agriculture (Hatch Project 1002996). The work conducted by the U.S. Department of Energy Joint Genome Institute, a DOE Office of Science User Facility, is supported by the Office of Science of the U.S. Department of Energy under Contract No. DE-AC02-05CH11231.

## FUNDING INFORMATION

U.S. NSF Microbial Observatories program (MCB-0702395) Katherine D McMahon

U.S. NSF Long Term Ecological Research program (NTL-LTER DEB-1440297) Katherine D McMahon

U.S. NSF INSPIRE award (DEB-1344254) Katherine D McMahon

U.S. National Institute of Food and Agriculture, U.S. Department of Agriculture (Hatch Project 1002996)

Katherine D McMahon

## LIST OF SUPPLEMENTARY MATERIAL

### Supplementary Text

#### Supplementary Figures S1 through S8

##### Supplementary Figure Legends

**Fig. S1.** A tiled display of an emergent self-organizing map (ESOM) based on the tetranucleotide frequency (TNF) of the 19 Verrucomicrobia MAGs. TNF was calculated with a window size of 5 kbp, with each dot on the ESOM representing a 5-kbp fragment (or a contig if its length is shorter than 5 kbp). Dots (i.e. fragments) are colored according to MAGs. A numeric ID is assigned to each MAG, and IDs from Mendota are labeled in black and IDs from Trout Bog labeled in white. A red outline was drawn to indicate the clustering of MAGs from Mendota on the ESOM.

**Fig. S2.** Counts of GH genes among the 78 different GH families present in MAGs.

**Fig. S3.** Heat map based on GH abundance profile patterns showing the clustering of MAGs by different lakes.

**Fig. S4.** Counts of carbohydrate and amino acid transporter genes.

**Fig. S5.** Comparison of glycolate oxidase gene operons in *E. coli*, *C. flavus* and TE4605.

**Fig. S6.** Summary of important metabolic genes and pathways.

**Fig. S7.** Occurrence and gene organization of Planctomycetes-specific domains, DUF1501, and DUF1552. (**a**) Counts of PSCyt, PSD, DUF1501, and DUF1552 domains in the MAGs. (**b**) Clustering of PUF1501- and PSCyt2-containing genes, and clustering of PUF1552- and PSCyt3-containing genes in the genome.

